# Prenatal Stress Differentially Shapes Adult Behavior in Male and Female Offspring

**DOI:** 10.64898/2026.02.16.705777

**Authors:** Erbo Dong, Alison Chu, Tamar Gur, Stephanie M. Gorka

## Abstract

**Background:** Prenatal stress (PNS) is a well-established risk factor for neuropsychiatric vulnerability, yet its sex-specific behavioral consequences remain incompletely defined. Because males and females follow distinct neurodevelopmental trajectories, clarifying how early-life stress differentially shapes behavior is essential for developing targeted interventions. However, few preclinical studies directly compare male and female offspring within the same experimental framework, limiting the ability to identify true sex-dependent effects.

**Methods:** Using a validated mouse model of gestational restraint stress, we conducted a comprehensive, within-study assessment of sex-dependent behavioral outcomes in adult offspring. Behavioral domains included locomotor activity, anxiety-like behavior, sociability, fear learning and extinction, recognition memory, and alcohol-related responses (ethanol preference and behavioral sensitivity), all measured using identical paradigms across sexes.

**Results:** PNS broadly disrupted behavior and cognition in both sexes, increasing locomotor activity and anxiety-like behavior, impairing fear extinction and recognition memory, and altering behavioral sensitivity to ethanol’s sedative effects. Direct comparison revealed distinct sex-dependent vulnerabilities: males showed reduced social interaction, whereas females exhibited numerically greater impairment in fear extinction and a significantly stronger ethanol preference. Baseline fear responses, total fluid intake, and sucrose consumption were unaffected.

**Conclusion:** Prenatal stress programs neurobehavioral trajectories in a sex-dependent manner, conferring vulnerability to anxiety-related behavior, cognitive disruption, and alcohol use. By directly comparing males and females within the same experimental design, this study provides one of the most integrated evaluations of sex-specific PNS outcomes to date and offers a robust framework for investigating the biological mechanisms underlying divergent pathways to stress-related psychopathology.

## INTRODUCTION

Early life represents a critical window of neurodevelopment during which environmental exposures can exert lasting effects on brain structure, function, and behavior (Miguel et al., 2020; Jubair, 2025; Peuters et al., 2024; Horton, 2024; Knudsen, 2004). Among these exposures, prenatal stress (PNS) is a particularly influential factor, associated with elevated risk for anxiety, depression, and substance use disorders later in life (Tung et al., 2023). Importantly, these outcomes are not uniformly expressed across sexes, with growing evidence that PNS shapes sex-specific trajectories of vulnerability (Georgousopoulou et al., 2025). Yet despite extensive human literature, preclinical studies rarely evaluate males and females side by side within the same experimental design, limiting the ability to distinguish true sex-dependent effects from methodological variability.

Clinical and naturalistic studies consistently demonstrate sex-specific responses to prenatal stress. Males often exhibit blunted cortisol reactivity, reduced placental buffering, and heightened sensitivity to early gestational stress, patterns linked to externalizing behaviors and increased risk for schizophrenia-spectrum and autism-like traits (Bale, 2011; Mueller & Bale, 2008). Females, by contrast, show heightened cortisol responses, compensatory placental adaptations, and greater sensitivity to later gestational stress, predisposing them to internalizing behaviors and mood-related disorders, including post-traumatic stress disorder (Kortesluoma et al., 2021; Sutherland & Brunwasser, 2018). Natural experiments such as the Dutch Hunger Winter and the Quebec Ice Storm further underscore these sex-specific outcomes: males show greater vulnerability to cognitive and psychotic disorders, whereas females exhibit higher rates of affective and emotional disturbances (Georgousopoulou et al., 2025; Levendosky et al., 2025). Beyond psychiatric risk, PNS is increasingly recognized as a determinant of sex-specific vulnerability to alcohol misuse, with males more likely to follow externalizing pathways and females more likely to follow stress-driven internalizing pathways that heighten risk for alcohol use disorder (Sutherland & Brunwasser, 2018; Levine et al., 2023). Despite these well-established sex differences in humans, preclinical PNS research has seldom incorporated alcohol-related outcomes in both sexes within the same study, leaving a critical translational gap. To address these limitations, we employed a mouse model in which offspring experience gestational restraint stress (Dong et al., 2014, 2018, 2022). Prior work from our group and others has characterized several behavioral consequences of this model, but most studies have focused on males or evaluated sexes separately, preventing direct comparison. The present study provides a comprehensive, within-study evaluation of male and female offspring using identical behavioral paradigms across locomotor activity, anxiety-like behavior, fear learning, social interaction, spatial learning, and alcohol-related responses. This design allows confirmatory findings to serve as essential anchors for interpreting sex differences and strengthens the model’s utility for mechanistic investigation and the development of sex-sensitive interventions.

These behavioral domains were selected because they reflect core functions disrupted by prenatal stress and are directly relevant to stress-related psychopathology. Prenatal stress is associated with anxiety, cognitive inflexibility, altered social behavior, and increased vulnerability to substance use (Weinstock, 2017; Van den Bergh et al., 2020). The chosen measures probe stress-sensitive neural circuits—including dopaminergic, amygdala–hippocampal, prefrontal–limbic, and mesolimbic pathways (Brunton, 2013; Bale, 2015)—and capture emotional, cognitive, and motivational consequences of early-life stress (Lupien et al., 2009). Assessing both sexes is essential for identifying sex-dependent neurobehavioral effects (Bale & Epperson, 2015; Gillies & McArthur, 2010). Together, these assessments advance the translational value of the PNS model and fill a critical gap by systematically defining sex-specific behavioral outcomes—including alcohol-related behavior—within a single, unified experimental framework.

## RESULTS

### PNS Produces Broad Behavioral Alterations, With Selective Sex-Specific Effects

Across behavioral domains, PNS produced robust alterations in locomotion, anxiety-like behavior, sociability, learning, memory, and alcohol-related responses. Several outcomes showed statistically confirmed sex × stress interactions, whereas others displayed numerical patterns consistent with sex-dependent vulnerability despite nonsignificant interactions. All statistical results are reported below.

### 1. PNS Increases Locomotor Activity Similarly in Males and Females

Open field testing revealed that both male and female PNS offspring (n = 10 per group) exhibited significantly elevated locomotor activity relative to NS controls (Figure 1).

**Figure 1.**
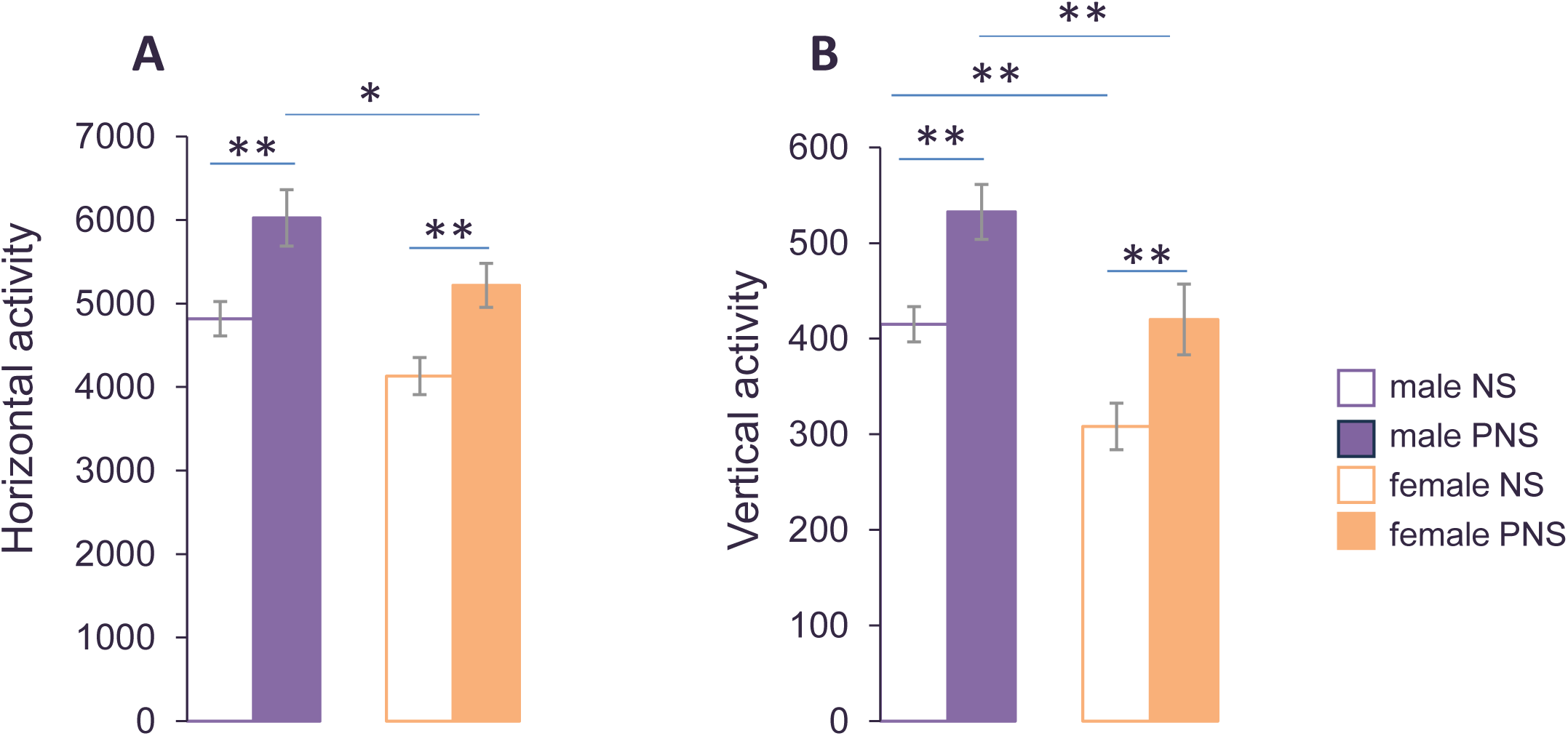
PNS increases locomotor activity in both sexes, with a numerical sex difference in vertical exploration. Male and female offspring exposed to PNS exhibited significantly elevated locomotor activity relative to NS controls. Horizontal activity increased similarly across sexes, whereas vertical activity showed a numerical but non-significant reduction in female PNS offspring compared with male PNS offspring. Data are presented as mean ± SEM (n = 10 mice per group). *p < 0.05; **p < 0.01.

For horizontal activity, two-way ANOVA showed significant main effects of stress, *F*(1, 36) = 19.151, *p* <.001, and sex, *F*(1, 36) = 8.125, *p* <.001, with no sex × stress interaction, *F*(1, 36) = 0.052, *p* =.821. Vertical activity showed a similar pattern, with significant main effects of stress, *F*(1, 36) = 16.817, *p* <.001, and sex, *F*(1, 36) = 15.384, *p* <.001, but no interaction, *F*(1, 36) = 0.10, *p* =.919. Female PNS offspring displayed numerically lower vertical activity than male PNS offspring, a pattern consistent with a potential sex-dependent alteration in rearing behavior, though the interaction was not significant.

### 2. PNS Heightens Anxiety-Like Behavior, With a Significant Sex-Specific Effect in the Light–Dark Box

#### Light–Dark Box (LDB)

In the LDB (n = 11 per group), PNS significantly increased time spent in the dark compartment for both sexes (Figure 2A). Two-way ANOVA revealed significant main effects of stress, *F*(1, 39) = 42.813, *p* <.001, and sex, *F*(1, 39) = 8.330, *p* =.006, as well as a significant sex × stress interaction, *F*(1, 39) = 4.223, *p* =.047. This interaction indicates a statistically confirmed sex-specific effect, with females showing a more pronounced dark-seeking response following PNS. Total ambulation did not differ across groups (stress: *F*(1, 39) = 2.877, *p* =.098; sex: *F*(1, 39) = 2.024, *p* =.163; interaction: *F*(1, 39) = 0.587, *p* =.448), confirming that anxiety-like behavior was not confounded by locomotor differences (Figure 2B).

**Figure 2.**
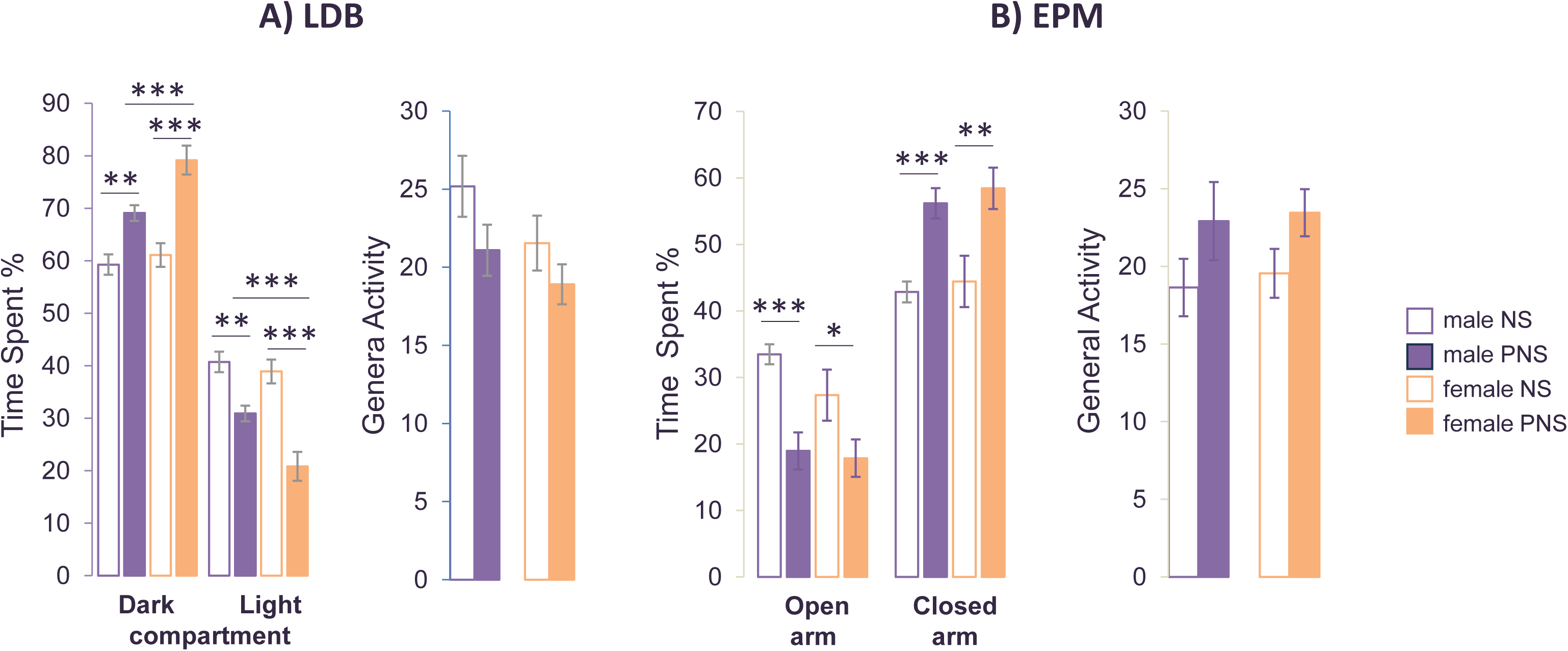
PNS heightens anxiety-like behavior, with a sex-specific effect emerging in the Light–Dark Box. Anxiety-related behavior was assessed using the Light–Dark Box (LDB) and Elevated Plus Maze (EPM). (A) In the LDB, PNS increased dark-compartment preference in both sexes, with females showing a significantly greater shift toward the dark compartment than males. (B) In the EPM, PNS reduced open-arm exploration in both sexes, with no significant sex × stress interaction. Data are presented as mean ± SEM (n = 11 mice per group). *p < 0.05; **p < 0.01; ***p < 0.001.

#### Elevated Plus Maze (EPM)

In the EPM, PNS reduced open-arm exploration in both sexes. For open-arm time, ANOVA showed a significant main effect of stress, *F*(1, 39) = 20.678, *p* <.001, but no effect of sex, *F*(1, 39) = 1.256, *p* =.269, and no interaction, *F*(1, 39) = 0.761, *p* =.388.

Closed-arm time showed a similar pattern (stress: *F*(1, 39) = 21.454, *p* <.001; sex: *F*(1, 39) = 0.126, *p* =.724; interaction: *F*(1, 39) = 0.490, *p* =.826). Total ambulation was unaffected (all *ps* >.08). Because the LDB and EPM tap partially distinct dimensions of threat and avoidance, these task-dependent effects likely reflect complementary, rather than inconsistent, aspects of anxiety-like behavior.

### 3. PNS Impairs Sociability in a Sex-Specific Manner

Sociability was assessed using the Three-Chamber Social Test (n = 10 per group). Two-way ANOVA revealed significant main effects of stress, *F*(1, 36) = 8.429, *p* =.006, and sex, *F*(1, 36) = 23.805, *p* <.001, as well as a significant sex × stress interaction, *F*(1, 36) = 5.326, *p* =.028.

Male PNS offspring exhibited significantly reduced social interaction compared to NS males (Figure 3). In contrast, PNS did not significantly alter sociability in females. Both NS and PNS females showed lower overall social engagement than males, reflecting a baseline sex difference in social investigation under these testing conditions, rather than an inference about social anxiety. This represents a statistically confirmed sex-specific effect, with males showing greater vulnerability to PNS-induced social deficits.

**Figure 3.**
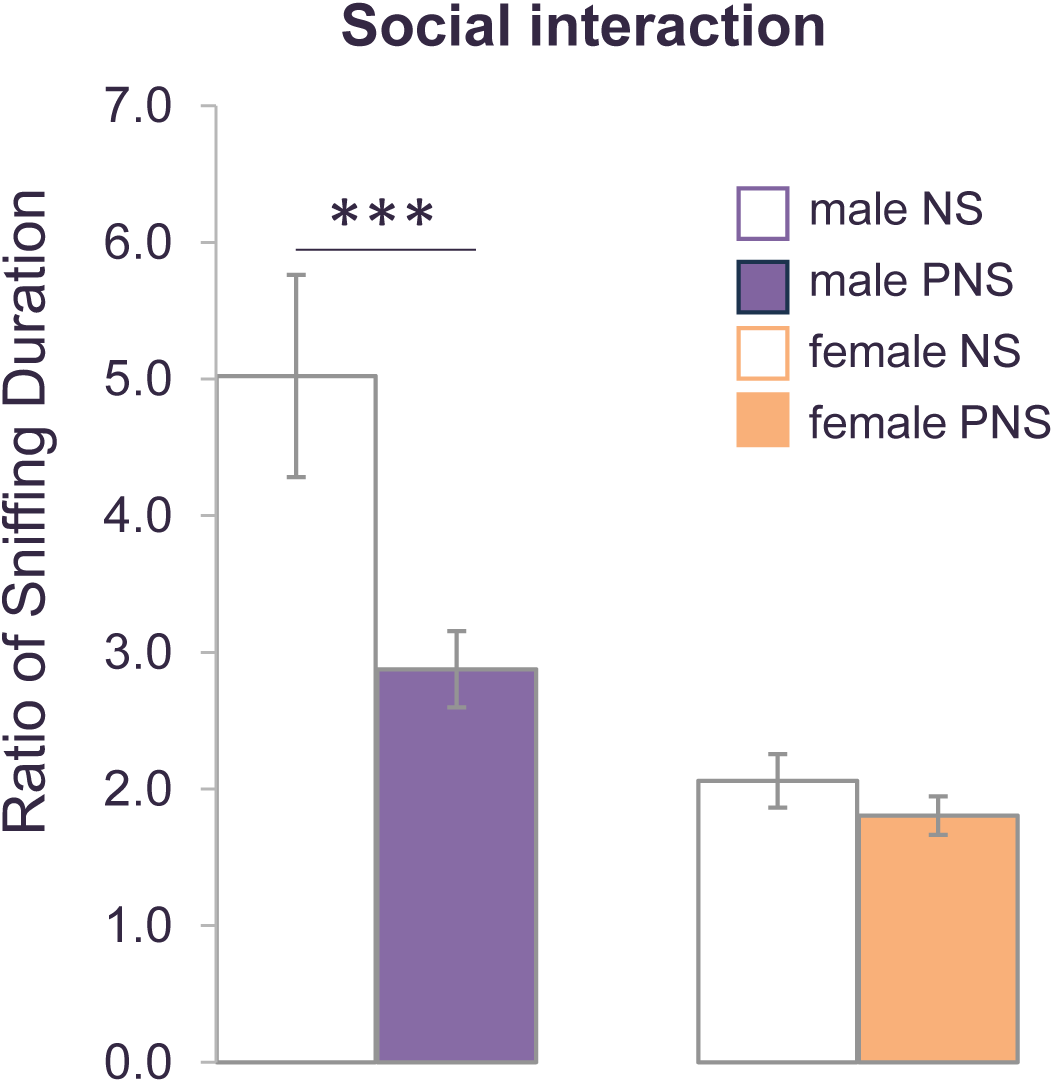
PNS reduces sociability in males but not females. Male PNS offspring displayed significantly reduced social interaction relative to NS males, whereas PNS did not significantly alter sociability in females. Both NS and PNS females showed lower baseline social investigation than males under these testing conditions. Social interaction was quantified as the ratio of time spent sniffing the stranger-mouse cup relative to the empty cup. Data are presented as mean ± SEM (n = 10 mice per group). ***p < 0.001.

### 4. PNS Impairs Fear Extinction, With Numerically Greater Impairment in Females

Fear extinction was examined in male and female NS and PNS offspring (n = 10 per group). Repeated-measures ANOVA revealed significant main effects of **stress**, *F*(1, 37) = 39.539, *p* <.001, and sex, *F*(1, 37) = 8.425, *p* =.006, but no sex × stress interaction, *F*(1, 37) = 0.453, *p* =.505. Both male and female PNS offspring showed impaired extinction relative to NS controls (Figure 4). Although females exhibited numerically greater impairment, this pattern did not reach significance and should be interpreted as a trend consistent with increased vulnerability, not a definitive sex-specific effect. Baseline freezing during habituation and conditioning did not differ across groups.

**Figure 4.**
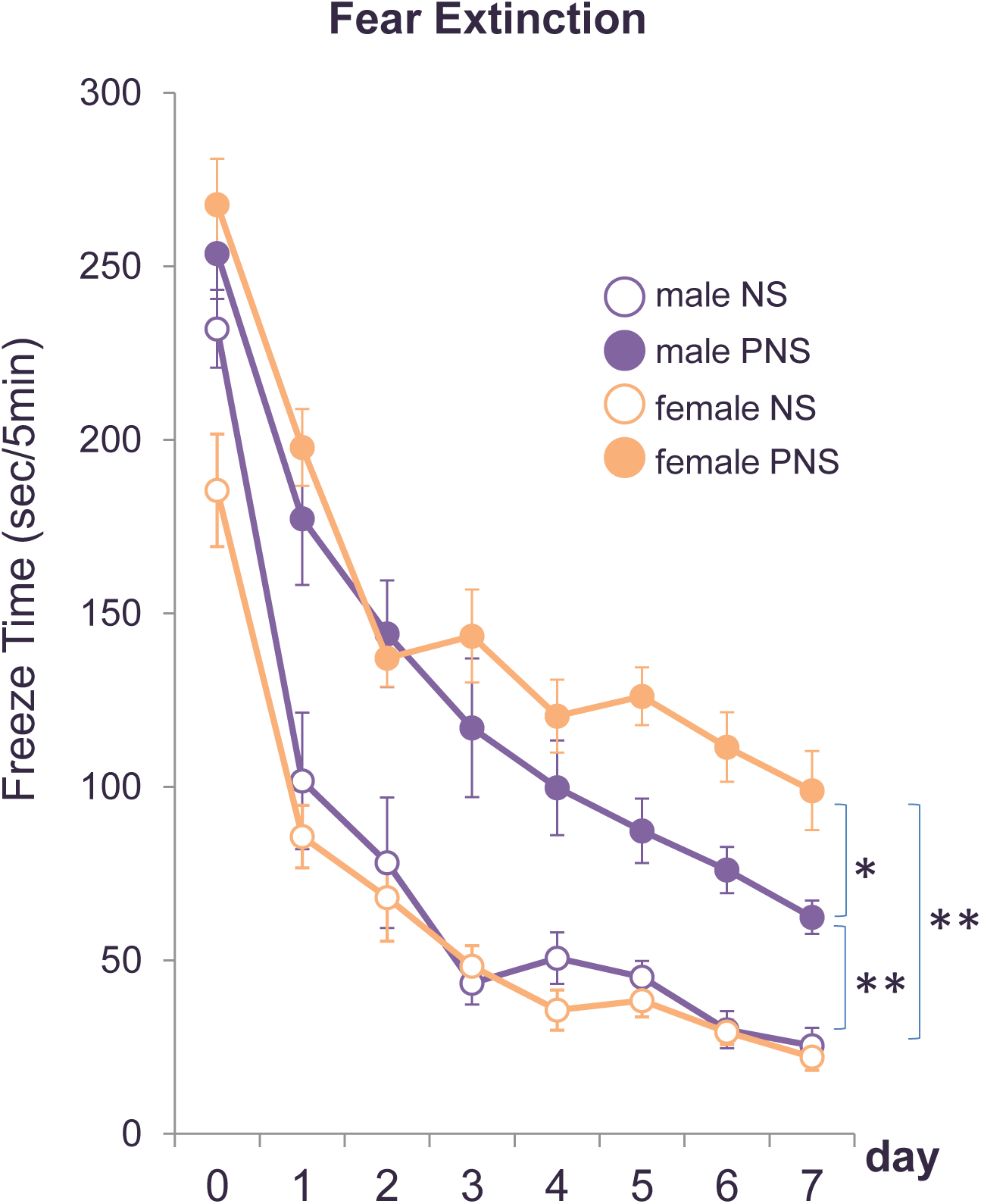
PNS impairs fear extinction in both sexes, with females showing numerically greater disruption. Following conditioning, NS offspring of both sexes exhibited robust freezing that declined across repeated context exposures, indicating extinction learning. PNS offspring showed significantly elevated freezing across extinction sessions, reflecting impaired extinction. Although females displayed numerically greater impairment, the sex × stress interaction was not statistically significant. Data are presented as mean ± SEM (n = 10 mice per group). *p < 0.05; **p < 0.01.

### 5. PNS Impairs Recognition Memory in Both Sexes

In the Novel Object Recognition (NOR) task (n = 11 per group), two-way ANOVA revealed a significant main effect of stress, *F*(1, 39) = 16.864, *p* <.001, but no effect of sex, *F*(1, 39) = 2.833, *p* =.100, and no sex × stress interaction, *F*(1, 39) = 0.235, *p* =.630.

Both male and female PNS offspring showed significantly reduced discrimination indices (Figure 5). Female PNS offspring displayed a numerically larger reduction, but this difference was not significant. Thus, PNS impaired recognition memory similarly across sexes.

**Figure 5.**
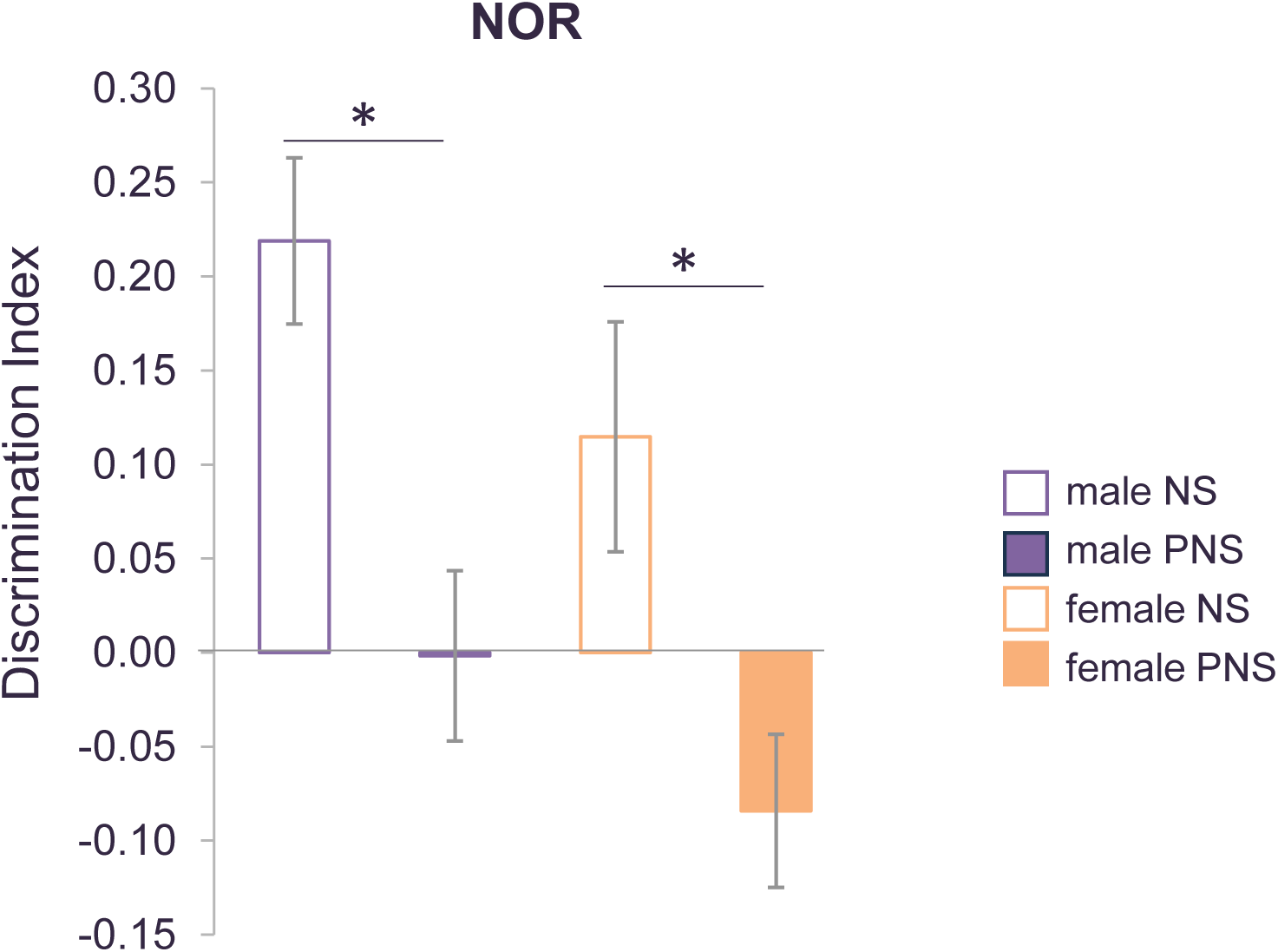
PNS impairs recognition memory in male and female offspring. Recognition memory was assessed using the Novel Object Recognition (NOR) test. During the test phase, both male and female PNS offspring showed significantly reduced discrimination indices (DI) relative to NS controls, indicating impaired ability to distinguish familiar from novel objects. Female PNS offspring exhibited a numerically larger DI reduction, though this difference was not statistically significant. Data are presented as mean ± SEM (n = 11 mice per group). *p < 0.05.

### 6. PNS Increases Ethanol Consumption, With a Significant Sex-Specific Increase in

#### Females

Voluntary ethanol consumption was assessed using a two-bottle free-choice paradigm (n = 11 per group). Two-way repeated-measures ANOVA revealed significant main effects of stress, *F*(1, 39) = 401.667, *p* <.001, and **sex**, *F*(1, 39) = 85.063, *p* <.001, as well as a significant sex × stress interaction, *F*(1, 39) = 56.099, *p* <.001. Female PNS offspring consumed substantially more ethanol than NS females and male PNS offspring across all concentrations (Figure 6). Follow-up analyses confirmed a trend toward a sex effect, *F*(1, 39) = 3.906, *p* =.055, a robust stress effect, *F*(1, 39) = 280.848, *p* <.001, and a significant sex × stress interaction, *F*(1, 39) = 5.712, *p* =.022. Total fluid intake did not differ across groups (stress: *F*(1, 39) = 0.601, *p* =.443; sex: *F*(1, 39) = 2.498, *p* =.122; interaction: *F*(1, 39) = 0.072, *p* =.790).

**Figure 6.**
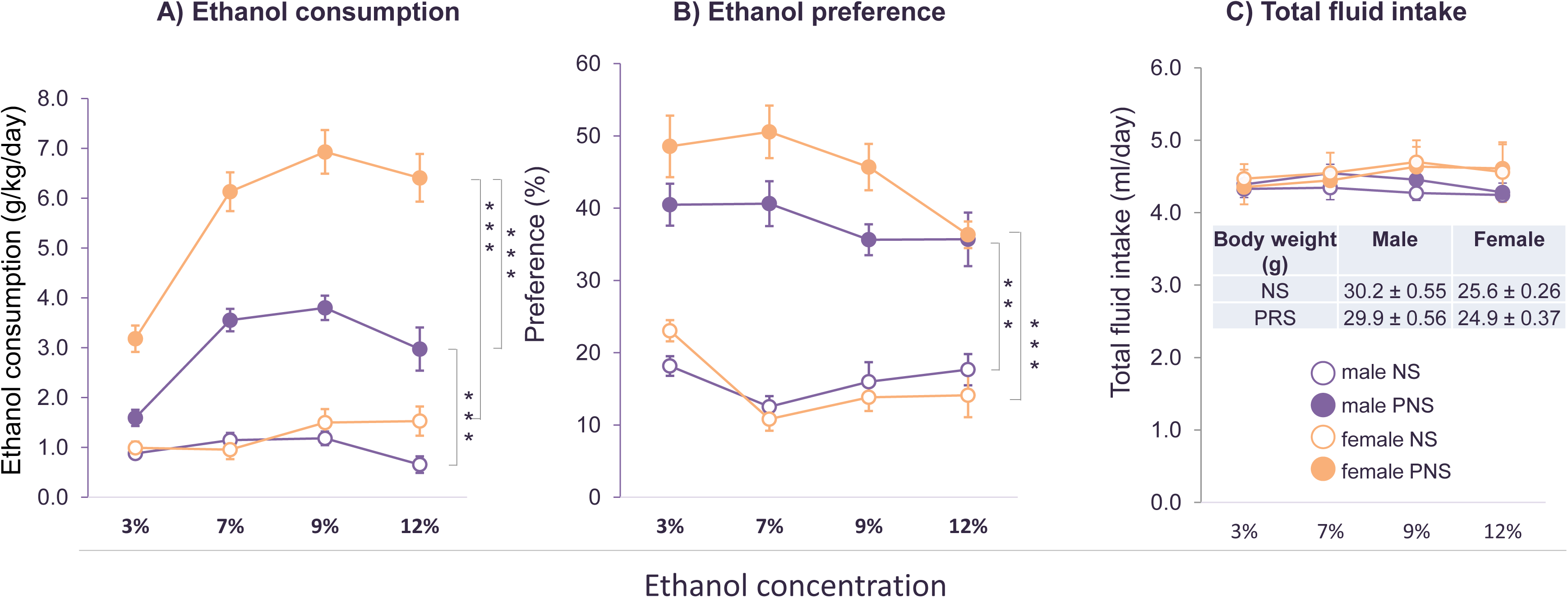
PNS increases ethanol consumption, with a significant sex-specific enhancement in females. (A) PNS significantly increased ethanol intake across concentrations in both sexes. (B) Water intake decreased correspondingly. (C) Total fluid intake and body weight did not differ across groups. Female PNS offspring consumed substantially more ethanol than male PNS offspring, reflecting a significant sex × stress interaction. Data are presented as mean ± SEM (n = 11 mice per group). ***p < 0.001.

Sucrose intake (n = 8 per group) showed no effect of stress, *F*(1, 28) = 0.284, *p* =.598, a significant effect of sex, *F*(1, 28) = 21.488, *p* <.001, and no interaction, *F*(1, 28) = 1.407, *p* =.246 (Figure 7). These findings demonstrate a statistically confirmed sex-specific increase in ethanol preference, with females showing the strongest intake.

**Figure 7.**
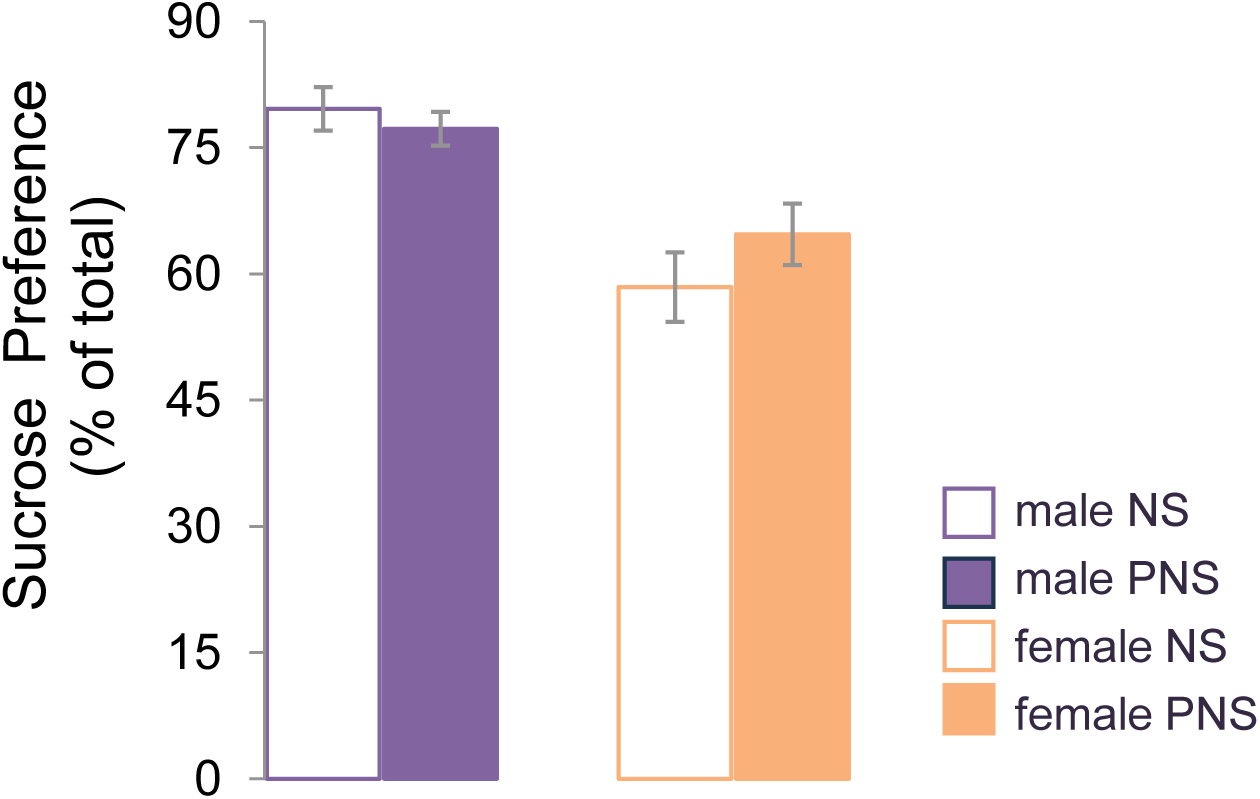
PNS does not alter sucrose preference. Sucrose preference did not differ between NS and PNS offspring. A significant main effect of sex indicated that females consumed less sucrose than males, independent of stress exposure. Data are presented as mean ± SEM (n = 8 mice per group).

### 7. PNS Alters Behavioral Sensitivity to Ethanol’s Sedative Effects

Ethanol sensitivity was assessed using loss of righting reflex (LORR) following 4.0 g/kg ethanol (n = 10 per group). For LORR latency, ANOVA revealed significant main effects of **stress**, *F*(1, 36) = 20.098, *p* <.001, and **sex**, *F*(1, 36) = 6.504, *p* =.015, but no interaction, *F*(1, 36) = 0.058, *p* =.810. For LORR duration, there was a significant main effect of stress, *F*(1, 36) = 26.742, *p* <.001, but no effect of sex, *F*(1, 36) = 0.414, *p* =.524, and no interaction, *F*(1, 36) = 0.483, *p* =.492. Both sexes showed prolonged latency and reduced duration following PNS (Figure 8), indicating altered behavioral sensitivity to ethanol’s sedative effects. Because blood ethanol levels were not measured, pharmacokinetic and pharmacodynamic contributions cannot be distinguished in the current study.

**Figure 8.**
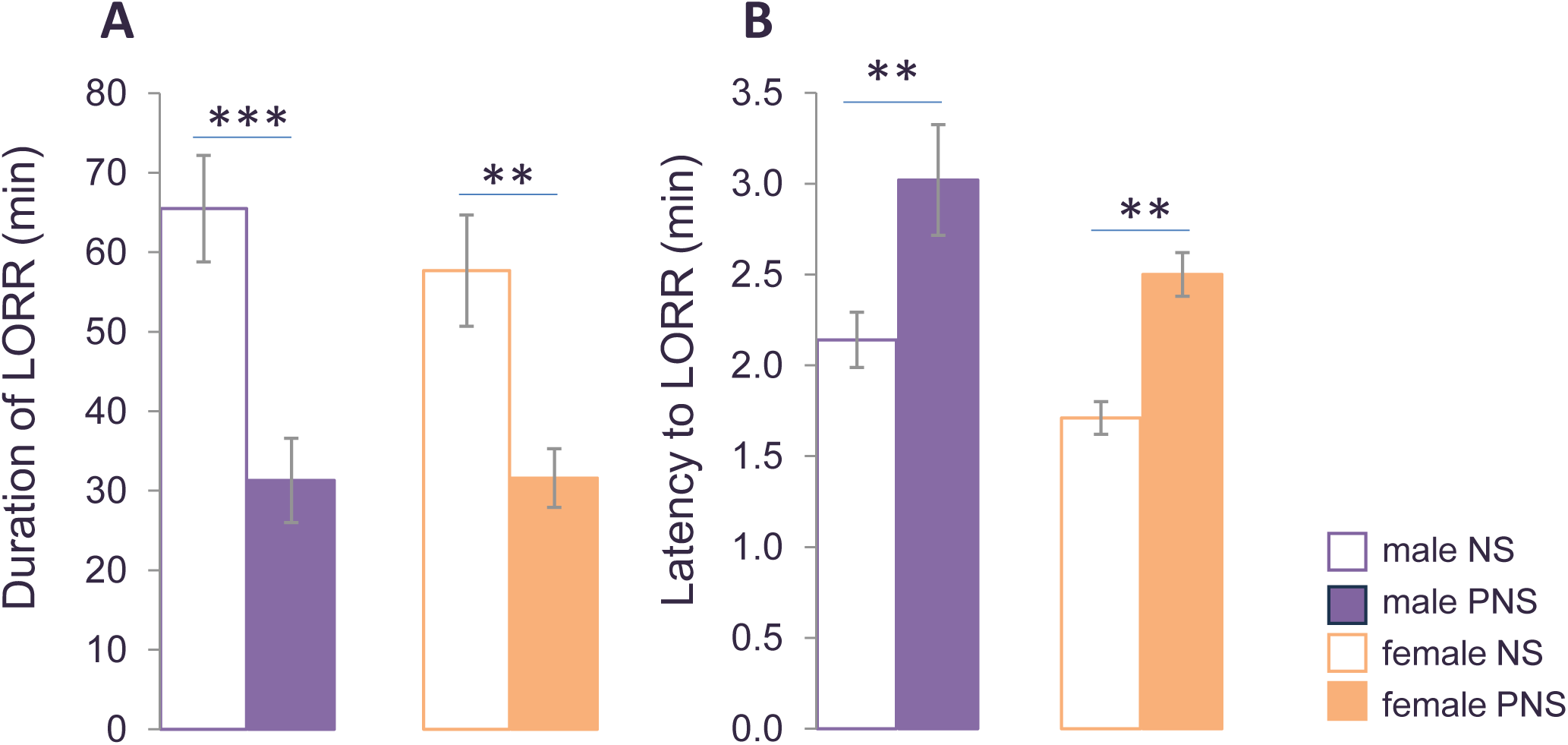
PNS alters behavioral sensitivity to ethanol’s sedative effects without sex-specific differences. Following 4.0 g/kg ethanol administration, PNS offspring of both sexes exhibited prolonged latency to loss of righting reflex (LORR) and reduced LORR duration relative to NS controls. No sex differences were detected. Because blood ethanol levels were not measured, pharmacokinetic versus pharmacodynamic contributions cannot be distinguished. Data are presented as mean ± SEM (n = 10 mice per group). **p < 0.01; ***p < 0.001.

## DISCUSSION

Our study demonstrates that PNS induces broad behavioral and neurocognitive alterations in adult offspring, with several outcomes showing clear sex-dependent effects. A major strength of this work is the direct, within-study comparison of males and females across multiple behavioral domains, which minimizes methodological variability and provides a more reliable foundation for interpreting sex differences.

Across assays, significant sex × stress interactions emerged in the light–dark box, social interaction, and ethanol preference, indicating robust sex-specific vulnerabilities in anxiety-related behavior, sociability, and alcohol-related motivation. In contrast, fear extinction and recognition memory showed numerically greater impairment in females, although interactions were not statistically significant. This pattern aligns with clinical observations of heightened internalizing vulnerability in females following early-life adversity, while respecting the statistical boundaries of the dataset.

In the anxiety domain, PNS increased avoidance behavior in both sexes, but the sex-specific interaction in the light–dark box suggests that females may be more sensitive to anxiogenic bright-light environments. The absence of a sex effect in the elevated plus maze is not contradictory; rather, these tasks probe partially distinct dimensions of threat and avoidance, with the light–dark box emphasizing light aversion and the elevated plus maze emphasizing open-space exposure. Together, these complementary assays indicate that PNS heightens anxiety-like behavior broadly, with sex-specific effects emerging under certain environmental conditions.

Social behavior showed a clear sex-specific deficit, with male PNS offspring exhibiting reduced social interaction, consistent with literature linking PNS to social withdrawal and autism-like traits in males. Female sociability was not significantly altered by PNS, and both NS and PNS females displayed lower baseline social investigation under these testing conditions, reflecting a sex difference in social exploration rather than an inference about social anxiety.

Cognitive outcomes were affected in both sexes. PNS impaired fear extinction and recognition memory, with females showing numerically larger reductions in extinction performance and discrimination index. Although these differences were not statistically significant, they are consistent with evidence that females may be more sensitive to stress-related disruptions in hippocampal-dependent cognition.

Alcohol-related behaviors revealed one of the strongest sex-specific effects. Female PNS offspring consumed substantially more ethanol than all other groups, reflecting a significant sex × stress interaction and suggesting that PNS may differentially shape motivational pathways related to alcohol use. Both sexes exhibited altered behavioral sensitivity to ethanol’s sedative effects, as indicated by prolonged latency and reduced duration of loss of righting reflex.

Because blood ethanol levels were not measured, pharmacokinetic versus pharmacodynamic contributions cannot be distinguished, and the findings should be interpreted as altered behavioral sensitivity rather than definitive changes in tolerance.

Together, these results reinforce the importance of treating sex as a biological variable in stress research. The within-study design clarifies where sex differences are statistically robust and where numerical trends suggest—but do not confirm—sex-specific vulnerability, providing a nuanced and statistically responsible interpretation of PNS effects.

### Potential Mechanisms Underlying Sex-Dependent Effects

Although mechanistic analyses were beyond the scope of the present study, several biologically plausible pathways may contribute to the observed sex-dependent outcomes:

#### Placental Function and Fetal Programming

Male and female placentas differ in glucocorticoid metabolism, inflammatory signaling, and growth factor expression. Reduced placental buffering in males and compensatory adaptations in females may shape sex-specific fetal exposure to maternal stress hormones.

#### Timing of HPA Axis Development

Critical windows of hypothalamic–pituitary–adrenal (HPA) axis maturation differ by sex. Stress during early gestation disproportionately affects males, whereas stress during later gestation more strongly impacts females, potentially contributing to the domain-specific patterns observed here.

#### Circuit-Level Vulnerabilities

PNS alters development of the amygdala, hippocampus, prefrontal cortex, and mesolimbic dopamine pathways. Sex differences in synaptic pruning, microglial activation, and excitatory–inhibitory balance may underlie divergent outcomes in anxiety, sociability, and alcohol-related behavior.

#### Hormonal and Neuroendocrine Factors

Sex hormones modulate stress responsivity, emotional regulation, and reward processing. Estrogen-and androgen-dependent signaling may amplify or buffer PNS effects in a domain-specific manner, particularly during adolescence and early adulthood.

#### Epigenetic Regulation

Prior work from our group has shown that PNS induces epigenetic alterations in genes regulating GABAergic and glutamatergic signaling in males. Extending these analyses to females will be essential for identifying sex-specific epigenetic signatures that may drive divergent behavioral trajectories.

## LIMITATIONS

While the PNS mouse model provides valuable insight into sex-dependent behavioral outcomes, several limitations should be considered. First, although this study offers a rare, within-study, sex-balanced comparison across multiple behavioral domains, rodent models cannot fully capture the complexity of human psychiatric conditions, which are shaped by species-specific differences in brain organization, genetics, and social context. Second, the restraint stress paradigm represents a well-defined and reproducible form of prenatal stress but does not encompass the diversity of stressors encountered by humans, including emotional trauma, socioeconomic adversity, or chronic environmental instability. Third, behavioral assays such as the Light–Dark Box and Elevated Plus Maze assess discrete components of anxiety-like behavior in simplified contexts and may not reflect the multidimensional nature of human anxiety. Fourth, sex hormone influences were not directly assessed, leaving open the possibility that hormonal fluctuations contributed to some of the observed sex differences. Finally, behavioral assessments were limited to late adolescence and early adulthood, preventing conclusions about lifespan trajectories, compensatory mechanisms, or the stability of sex differences across development.

These limitations highlight the need for complementary approaches—including longitudinal human studies, diverse stress paradigms, and molecular and endocrine analyses—to deepen our understanding of sex-specific vulnerability to early-life stress. Importantly, they do not diminish the central strength of this work: a controlled, within-study comparison of male and female offspring using identical behavioral paradigms, providing a robust foundation for interpreting sex-dependent effects of prenatal stress.

## CONCLUSION

This study demonstrates that prenatal stress exerts broad and partially sex-dependent effects on behavioral and cognitive outcomes relevant to psychiatric and substance-use vulnerability.

Significant sex × stress interactions emerged in anxiety-related behavior, sociability, and ethanol preference, while other domains—such as fear extinction and recognition memory—showed numerical patterns consistent with greater female vulnerability, though without statistically significant interactions. Both sexes exhibited altered behavioral sensitivity to ethanol’s sedative effects and impairments in recognition memory, indicating shared vulnerabilities alongside domain-specific divergences.

A key strength of this work is its direct, within-study comparison of males and females across multiple behavioral domains, allowing confirmatory findings to serve as anchors for interpreting sex differences and providing a clearer, more reliable picture of how PNS shapes neurobehavioral trajectories. By integrating internalizing, externalizing, cognitive, and substance-related phenotypes within the same cohort, this study offers one of the most comprehensive sex-balanced behavioral characterizations of PNS to date.

These findings support the hypothesis that early-life stress programs neurobehavioral outcomes in a sex-dependent manner, potentially through distinct molecular, hormonal, placental, and circuit-level pathways. The PNS model thus provides a powerful platform for investigating the biological mechanisms underlying stress-related disorders and for developing targeted, sex-specific interventions. Understanding how prenatal stress shapes vulnerability to psychiatric conditions is essential for informing public health strategies focused on maternal care and early-life prevention. Future research integrating neuroendocrine profiling, epigenetic mapping, and cross-species comparisons will be critical for bridging the translational gap and enhancing therapeutic precision.

## METHODS

### Animals and PNS Procedure

All procedures followed NIH guidelines for the Care and Use of Laboratory Animals (NRC, 1996) and were approved by the University of Illinois at Chicago Animal Care Committee. Pregnant Swiss albino ND4 mice (Harlan, Indianapolis, IN, USA) were individually housed under a 12-h light–dark cycle with ad libitum access to food and water. Control dams (NS) were left undisturbed throughout gestation. Stressed dams (PNS) underwent a repeated restraint stress paradigm as previously described (Dong et al., 2014, 2016; Zheng et al., 2016; Matrisciano et al., 2013). Beginning on gestational day 7 and continuing until delivery, dams were placed in a transparent Plexiglas tube (12 × 3 cm) under bright light for 45 minutes, three times daily. After weaning on postnatal day 21, male and female offspring were group-housed (five per cage) by condition. To minimize litter-based confounds, no more than one or two pups of each sex from a given litter were assigned to any behavioral measure (Becker & Kowall, 1977; Chapman & Stern, 1979). All behavioral tests—including drinking experiments—were conducted in the same cohort of mice.

The estrous cycle was not monitored. Although many stress-related behavioral phenotypes are robust across estrous stages, the absence of estrous tracking is acknowledged as a methodological limitation.

### Locomotor Activity

Locomotor activity was assessed using a computerized Animal Activity Monitoring System with VersaMax software (AccuScan Instruments, Columbus, OH), following established procedures (Dong et al., 2014). Each chamber consisted of a transparent Perspex box (20 × 20 × 20 cm) surrounded by horizontal and vertical infrared beams. Horizontal activity was quantified by interruptions of horizontal beams. Vertical activity (rearing) was quantified by interruptions of vertical beams. Each mouse was tested for 15 minutes between 1:00 PM and 3:00 PM. Data were collected automatically and analyzed offline.

### Light/Dark Box (LDB) Exploration Test

Anxiety-like behavior and general locomotor activity were assessed using the Light/Dark Box (Dong et al., 2018). The apparatus consisted of two connected compartments: a dark chamber with no illumination, and a light chamber illuminated by a 0.25-Amp LED.

Mice were acclimated to the testing room for 5 minutes before testing. Each mouse was placed into the dark compartment facing away from the opening and allowed to explore freely for 5 minutes. Behavior was recorded using an automated tracking system.

For each animal, we quantified: time spent in the light and dark compartments, percentage of time in each compartment, and total ambulation across both chambers as an index of general activity.

### Elevated Plus Maze (EPM)

Anxiety-like behavior was also assessed using the Elevated Plus Maze (Dong et al., 2018). The apparatus consisted of two open arms and two closed arms constructed from Plexiglas and elevated 50 cm above the floor. Mice were placed on the central platform facing an open arm and allowed to explore for 10 minutes. Behavior was recorded using an automated tracking system.

The primary measure was percentage of time spent in open arms, calculated as:

(open-arm time × 100) / (open-arm time + closed-arm time). Closed-arm entries served as an index of general locomotor activity.

### Three-Chamber Social Interaction Test

Social approach behavior was assessed using a transparent Plexiglas three-chamber apparatus (Dong et al., 2014). Each chamber measured 20 × 40.5 × 22 cm and was connected by small openings allowing free movement. Two identical wire cups were placed in the side chambers—one empty and one containing a novel conspecific. After a 5-minute habituation period, the test mouse was allowed to explore all chambers for 10 minutes. Social interaction was quantified as the ratio of time spent sniffing the stranger-mouse cup relative to the empty cup. Reliability was confirmed by independent scoring from two trained raters. The apparatus was cleaned with 70% ethanol and distilled water between trials.

### Novel Object Recognition (NOR)

NOR testing followed established procedures (Lueptow, 2017) and occurred over three days: Day 1: Habituation; Day 2: Training with two identical objects; Day 3: Testing with one familiar and one novel object. Each session lasted 10 minutes. To ensure rigor: A third copy of each object was used during testing to avoid olfactory cues. Object identity and location were counterbalanced across mice. The diagonal used during training was matched during testing.

The discrimination index (DI) was calculated as:

(novel exploration − familiar exploration) / total exploration.

Exploration was defined as nose-oriented investigation within 2–3 cm. All apparatus and objects were cleaned with 70% ethanol and distilled water between trials.

### Alcohol Preference Test

Voluntary ethanol consumption was assessed using a two-bottle free-choice paradigm (Dong et al., 2018, 2022). Individually housed mice received ad libitum access to water and ethanol solutions. Bottle positions were alternated daily to prevent side bias. After water habituation, mice received ethanol concentrations of 3%, 7%, 9%, and 12% (v/v), each for three days. Fresh solutions were provided daily at the onset of the dark cycle. Fluid intake was recorded at 6:00 PM. Ethanol preference was calculated as the percentage of ethanol intake relative to total fluid intake. Ethanol consumption was expressed as g/kg/day.

### Sucrose Preference Test

Sucrose preference was assessed using the same two-bottle free-choice paradigm (Dong et al., 2022), with mice individually housed. After water habituation, mice received 0.2%, 2%, and 5% sucrose solutions, each for two days. Bottle positions were alternated daily. Fluid consumption was measured at 6:00 PM. Sucrose intake was expressed as g/kg/day, and sucrose preference was calculated as the ratio of sucrose intake to total fluid intake.

### Fear Conditioning and Extinction

Fear conditioning was conducted in a transparent acrylic chamber with a grid floor capable of delivering a mild foot shock (2 s, 0.5 mA; San Diego Instruments). Mice received three pairings of a 30-s, 85-dB tone (CS) co-terminating with a foot shock (US) during a 2-minute training session. Twenty-four hours later, contextual fear memory was assessed by returning mice to the same chamber for 5 minutes without tone or shock. Contextual testing was repeated daily for up to seven days to assess extinction. Freezing was defined as the absence of movement except respiration. Cued fear memory was assessed in a novel chamber with distinct visual cues, where the tone was presented without shock.

### Loss of Righting Reflex (LORR)

Behavioral sensitivity to ethanol’s sedative effects was assessed using LORR (Pardo et al., 2013). Mice received an intraperitoneal injection of 4.0 g/kg ethanol and were placed individually in a clean cage. Latency to LORR: time from injection to loss of righting reflex Duration of LORR: time from onset of LORR to recovery, defined as three successful rightings within one minute. Mice were positioned on their backs in a V-shaped cradle to prevent spontaneous righting. Testing occurred in a quiet room with soft lighting. Because blood ethanol levels were not measured, pharmacokinetic and pharmacodynamic contributions to LORR cannot be distinguished.

## ACKNOWLEDGMENTS

This study was supported by the NIH-NIAAA grant R21AA027848 to ED

## CONFLICT OF INTEREST

The authors declare no competing interests.

## Notes

### Competing Interest Statement

The authors have declared no competing interest.

